# Joint disruption of *Ret* and *Ednrb* transcription drives cell fate reversal in the Enteric Nervous System in Hirschsprung disease

**DOI:** 10.1101/2025.03.04.641483

**Authors:** Ryan D. Fine, Rebecca Chubaryov, Mingzhou Fu, Gabriel Grullon, Aravinda Chakravarti

**Affiliations:** Center for Human Genetics and Genomics, NYU Grossman School of Medicine, New York, NY; Department of Neuroscience and Physiology, NYU Grossman School of Medicine, New York, NY

**Keywords:** *Ret*, *Ednrb*, aganglionosis, gene expression, mouse

## Abstract

Despite extensive genetic heterogeneity, 72% of pathogenic alleles for Hirschsprung disease (HSCR) arise from coding and regulatory variants in genes of the *RET* and *EDNRB* gene regulatory network (GRN) in the enteric nervous system (ENS). Reduced signaling of these two receptors below a threshold in enteric neural crest-derived cells (ENCDCs) leads to a molecular tipping point at which otherwise lesser cellular defects result in aganglionosis. To elucidate the mechanisms leading to enteric neuronal loss from these genetic defects, we generated four strains of mice carrying reduced function alleles at *Ret* or *Ednrb* or both, along with their wildtype alleles. ENS tissue- and single-cell gene expression profiling of the developing and postnatal gastrointestinal tract in five mouse models, with various combinations of mutant alleles, revealed 3 major insights: (i) *Ret* and *Ednrb* deficiency, rather than complete loss, is sufficient to induce HSCR, (ii) *Ret* and *Ednrb* demonstrate strong trans interactions, and (3) disruption of this interaction leads to cellular fate changes to compensate for neuronal loss. This study of targeted mouse models of a multifactorial disorder reveals how increasing dosage of genetic defects within a GRN leads to quantifiably increasing dysregulation from genotype to gene expression to cellular identity to function. Importantly, our studies establish that aganglionosis results only with severely reduced gene expression at both receptor genes and their consequent disruption of normal and compensatory cell fate trajectories.

## Introduction

The majority of human traits and disorders are multifactorial. The genetic component of such traits comprises both coding genes as well as their numerous cis-regulatory elements (CREs): sequence variation within both classes of genetic elements collectively shape the transcriptional and translational output of specific cells to affect a phenotype and its variation between individuals.^1,2^ Genomic studies of complex traits have mapped these multifactorial effects to hundreds or even thousands of genes,^3,4^ but the route by which these genetic elements modulate a phenotype is unknown. Pritchard and colleagues have argued that complex traits are likely ‘omnigenic’ with many mapped genes being trait-unrelated passengers whose expression is altered simply by being in the same regulatory network as the many fewer core genes.^5,6^ Nevertheless, the key question remains: how do the many genes of a trait modulate its phenotype? How does the responsible cell-type(s) know that it has 3 and not 30 variants, particularly when the variants are in diverse genes? How does the genetic information across an individual’s genome get integrated into a specific phenotype? Over the past decade, we have advanced a specific hypothesis to explain this enigma: trait genes cluster within one or more gene regulatory networks (GRNs) to regulate the expression of a few rate-limiting core genes that regulate an essential cell phenotype (e.g., differentiation).^2^ It is the quantitative difference of 3 or 30 variants on the rate-limiting core genes that alters the cell phenotype and, thereby, the organismal phenotype.

The GRN hypothesis emerged from our many studies of the enteric neuro-developmental disorder Hirschsprung disease (HSCR), an almost exclusively congenital multifactorial disease of the enteric nervous system (ENS) with megacolon and varying segments of cranio-caudal aganglionosis.^2–4^ In HSCR, a core set of 10 genes comprise the GRN that regulates the expression of two key receptor genes, *RET* and *EDNRB,*^5,6^ controlling the proliferation and migration of enteric neural crest-derived cells (ENCDCs).^7^ The dysregulation of this last step leads to the absence of intestinal ganglia in the submucosal plexuses from the dearth of cells necessary to populate the gut and form a complete ENS.^8^ This model explains some genetic features of HSCR, such as the overwhelming contribution of the *RET*-*EDNRB* GRN to HSCR attributable risk,^9^ the dose-dependent effects of disease variants on *RET* and *EDNRB* gene expression,^10^ how disease variants affect fundamental cell phenotypes (proliferation and migration) and why hundreds to thousands of genes are affected.^7^ However, the model does not reveal the mechanism of the dosage dependence on clinical disease: how much GRN dysregulation and how much loss of proliferation and migration is necessary to precipitate aganglionosis? To do so, we generated and studied specific mouse strains to model *Ret* and *Ednrb* dysregulation through differential loss of their gene expression.

*RET* and *EDNRB* are the major sites of susceptibility variants in HSCR. It has been classically assumed that loss-of-function (LoF) of these genes is necessary to cause aganglionosis, based on mouse models with homozygous null alleles.^11–13^ However, these mice fail to demonstrate two significant features of HSCR, namely, a higher male penetrance (incidence) and variation in the length of aganglionosis. To create more representative disease models, we previously generated compound mutant mouse strains with a null mutation at *Ret* and both null and hypomorphic mutations at *Ednrb*.^14,15^ These studies demonstrated the expected male sex-bias in aganglionosis and segment length variation, as well as a gut-specific genetic interaction between *Ret* and *Ednrb*. In later studies, we explored the gene expression consequences of such mutations,^16,17^ *Ret* null mutations in the developing mouse gut,^8,18^ and *RET* enhancer variants in human embryonic gut.^19^ These studies revealed the widespread cell autonomous and non-cell autonomous consequences of reduced function with *Ret* deficiency reducing ENCDC numbers and increasing (relatively) non-ENCDC cell numbers. However, the precise role of the *Ret*-*Ednrb* interaction in regulating penetrance remained elusive.

In this study we address this penetrance (incidence) question by conducting tissue-level and single-cell gene expression studies of the developing (embryonic day E14.5) and postnatal (P0) gastrointestinal tract in five mouse strains, with various combinations of mutant alleles at *Ret* or *Ednrb* or both, in contrast to the WT. We demonstrate that epistasis between *Ret* and *Ednrb* deficiency alleles dysregulates a specific set of 225 genes that neither single gene deficiency genotype does alone or do so additively. These gene sets have a highly specific pattern of expression dynamics with joint deficiency allele dosage and, we demonstrate, that this pattern arises from a shift in cell fate from epithelial/mesenchymal to neuronal, providing evidence for the first time that HSCR involves the loss of cellular function due to premature cell fate transitions beyond its known cell proliferation and migration defects.

## Results

### Ret and Ednrb mouse models used

In earlier work, our group reported on the consequences of a gene-dosage series of the *Ednrb* piebald and piebald lethal mutations, *Ednrb^s^* and *Ednrb^sl^*, crossed to a knockout allele of *Ret.*^14^ While *Ret* and *Ednrb^sl^* mutations are traditional null alleles, the piebald mutation has been reported to reduce *Ednrb* transcription in homozygotes by 75% due to alternative splicing of a 5.5kb retrotransposable element inserted in intron 1.^20^ Analysis of these mutations suggested that ∼50% expression of each gene was the threshold between enteric ganglionosis and aganglionosis. More importantly, the *Ret^+/-^ Ednrb^s/s^* model exhibited human-like sex bias in length of aganglionic tissue.^15^ One drawback of these mouse models was the mixed genetic backgrounds, primarily of 129S1/SvIMJ origin, leaving open the possibility that some of the phenotypic variation arose from the strain genetic background, as seen in the sex-bias penetrance of an *Ednrb^sl^* rat model.^21^ Thus, we began by acquiring two strains in an isogenic C57Bl6/J background. While the piebald mutation remains unchanged from our earlier work, the *Ret* null mutation used in this study is a cyan fluorescent protein (CFP) knockin-knockout model in which the *CFP* open reading frame, coupled to a poly-A tail, was inserted into exon 1 of *Ret* to disrupt function while allowing visualization of *Ret*-positive enteric neurons throughout development.^13^ We did not include heterozygous and homozygous *Ret-CFP* alleles in the majority of analyses for two reasons: First, this would require separate breeding and thus separate sample handling, and second, we did not want to re-model the *Ret* homozygous null as it is an outlier with complete loss of enteric neurons.^13^ However, we did include the *Ret* WT as the baseline control (**Figure 1A**).

**Figure 1.**
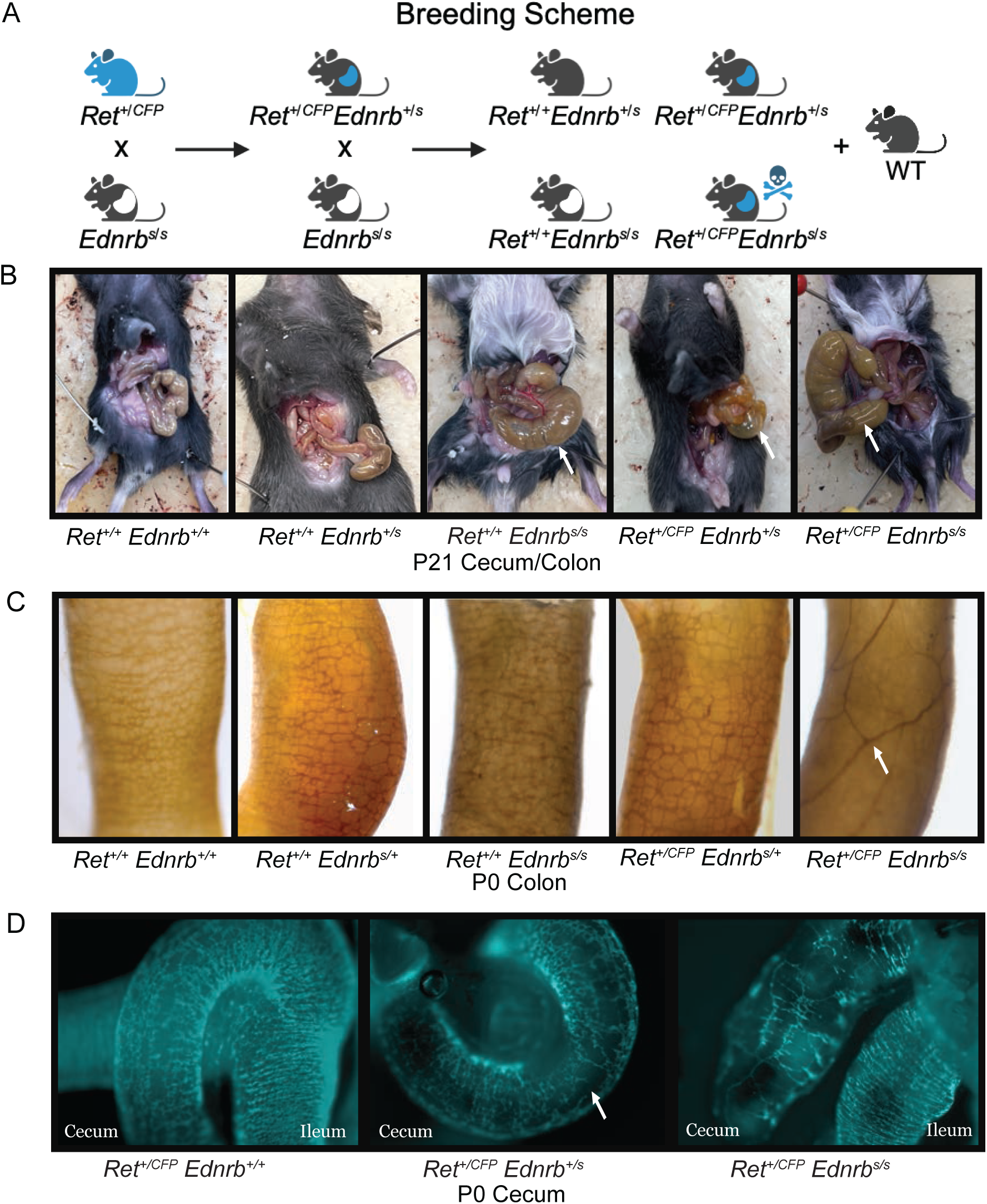
Phenotyping a series of increasingly HSCR-sensitive mouse lines. **(A)** Breeding scheme for four mouse mutant genotypes, excluding WT breeding. **(B)** Anatomical images of mouse colon relative to body size at P21. Panels 1 & 2: Cecum of a phenotyptically normal mouse. Panel 3: Partial megacolon in the cecum of a male mouse euthanized for sluggish movement. Panel 4: Translucent cecum of a male mouse euthanized for sluggish movement. Panel 5: A mouse with severe megacolon. **(C)** Representative Acetylcholinesterase (AChE) staining of the distal mouse colon at P0 for all genotypes, arranged in order from the least to the greatest risk of aganglionosis. White arrow highlights elongated and sparse neurons, a characteristic of the human HSCR gut. The frequency is quantified in **Table 1**. **(D)** Fluorescent images of *Ret* promoter-driven CFP expression in the whole-mount cecum of P0 animals in genotypes from the least to the greatest risk of aganglionosis. Left: Functionally dense neuronal network. Middle: Arrow highlights region with sparse/missing neurons in a phenotypically WT animal. Right: Full-blown aganglionosis in the cecum but not the ileum of a mutant mouse.

**Table 1.**
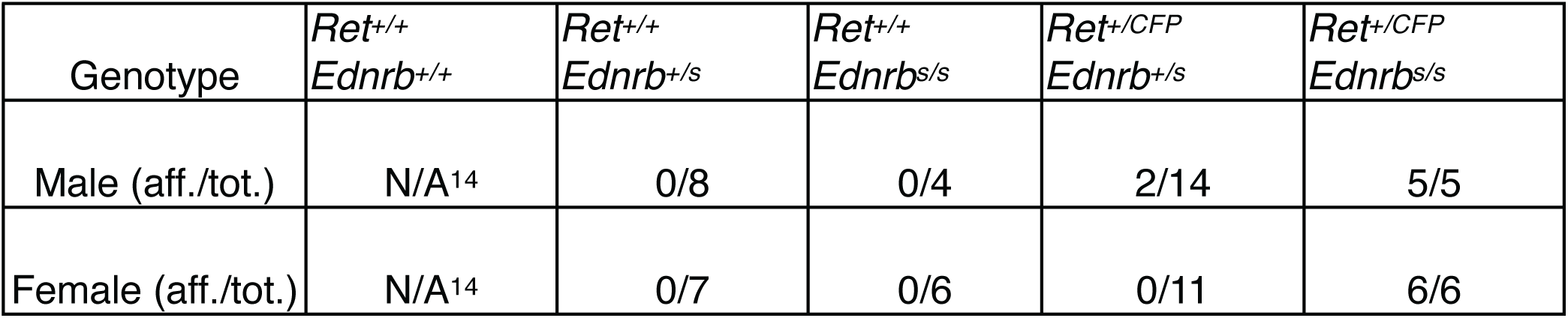
Aganglionosis Penetrance. Intestine from mice of the represented genotypes was dissected out at P0 and quantified by AChE staining for presence of nerve fibers. Mice scored as affected exhibited hypoganglionosis in the colon at minimum. Wildtype mice are assumed to be unaffected as previously reported.^14^

### Ret and Ednrb deficiency rather than absence is sufficient to cause *HSCR*

We allowed several litters of mice to reach maturity during initial colony expansion and breeding to assess overall health. *Ret^+/CFP^ Ednrb^s/s^* mice were able to survive up to P21, however, they were all eventually lost to HSCR-induced defects, notably abdominal distention from megacolon, one of the hallmarks of HSCR arising from an extensive swelling of gut tissues due to the inability to pass stool (**Figure 1B**). Additionally, we qualitatively observed several instances of sluggishness among the *Ret^+/CFP^ Ednrb^+/s^* and *Ret^+/+^ Ednrb^s/s^* male mice in which postmortem dissection revealed a translucent cecum or partial megacolon in the cecum, respectively (**Figure 1B**). Indeed, the cecum presents a significant obstacle for migrating enteric neurons with its unusual three-dimensional structure, since neurons must migrate both along the intestine and transmigrate along the mesentery to fully colonize the colon. Note that the cecum is the canonical landmark used to distinguish the colon from the small intestine.

We wanted to assess penetrance quantitatively, so we examined 61 pups across four mutant genotypes at postnatal day 0 (P0) by gross anatomical dissection and histochemical acetylcholinesterase (AChE) staining of the colonic submucosa to examine ENS neuronal ganglia.^16^ Our results showed 100% penetrance of the loss of AChE positive enteric neurons in the colons of *Ret^+/CFP^ Ednrb^s/s^* mice, as previously demonstrated (**Table 1**, **Figure 1B**).^14^ We noticed that a small (2/14) number of *Ret^+/CFP^ Ednrb^+/s^* male mice also exhibited mild loss of enteric neurons in the P0 colon. Using fluorescent microscopy, we examined 6 *Ret^+/CFP^ Ednrb^+/s^* mice and found that 50% (2 males, 1 female) of these exhibited mild neuronal loss in the cecum (**Figure 1D, middle panel**). This phenotype also appeared more severely in 31% (4/13) of female *Ret^+/CFP^ Ednrb^s/s^* mice (**Figure 1D**, **right panel**), while the remainder exhibited total loss of *CFP-*positive neurons in the cecum. Collectively, this demonstrates that the *Ret-CFP* and *Ednrb-s* allelic series is a better model for the human disease than classical homozygous knockout models, since they all have wildtype levels of neuronal density in the small intestine that is stochastically depleted from the cecum to the anal pore even in the most severe *Ret^+/CFP^ Ednrb^s/s^* genotype (**Figure S1**).

### Gene expression changes across developmental time, sex, and genotype

The four mutant strains we generated differed in their *Ret and Ednrb* deficiencies as well as their disease penetrance (aganglionosis); among these, only *Ret^+/CFP^ Ednrb^s/s^* have 100% disease penetrance. To identify the molecular genetic differences that produce complete from low penetrance with small genetic changes, suggesting a non-additive interaction, we performed bulk RNA-seq followed by statistical analysis of interactions after accounting for *Ret* and *Ednrb* additive allele dosage effects and sex. The gene expression data were quantified as absolute read counts and normalized for library size. To cover the many sources of variation, we studied 5 genotypes (WT plus 4 mutants of increasing severity: *Ret^+/+^ Ednrb^+/+^*, *Ret^+/+^ Ednrb^+/s^*, *Ret^+/+^ Ednrb^s/s^*, *Ret^+/CFP^ Ednrb^+/s^*, *Ret^+/CFP^ Ednrb^s/s^*), both sexes, two timepoints and biological triplicates for a total of 60 unique samples (**Figure 2A**). In addition to the P0 colon, we included the entire intestinal tract from E14.5, by which time the ENS is established.^22^ For analysis we utilized a general linear model in which the average gene expression of each *Ret* and *Ednrb* genotype was used to eliminate genes whose expression could be explained through their simple additive action.

**Figure 2.**
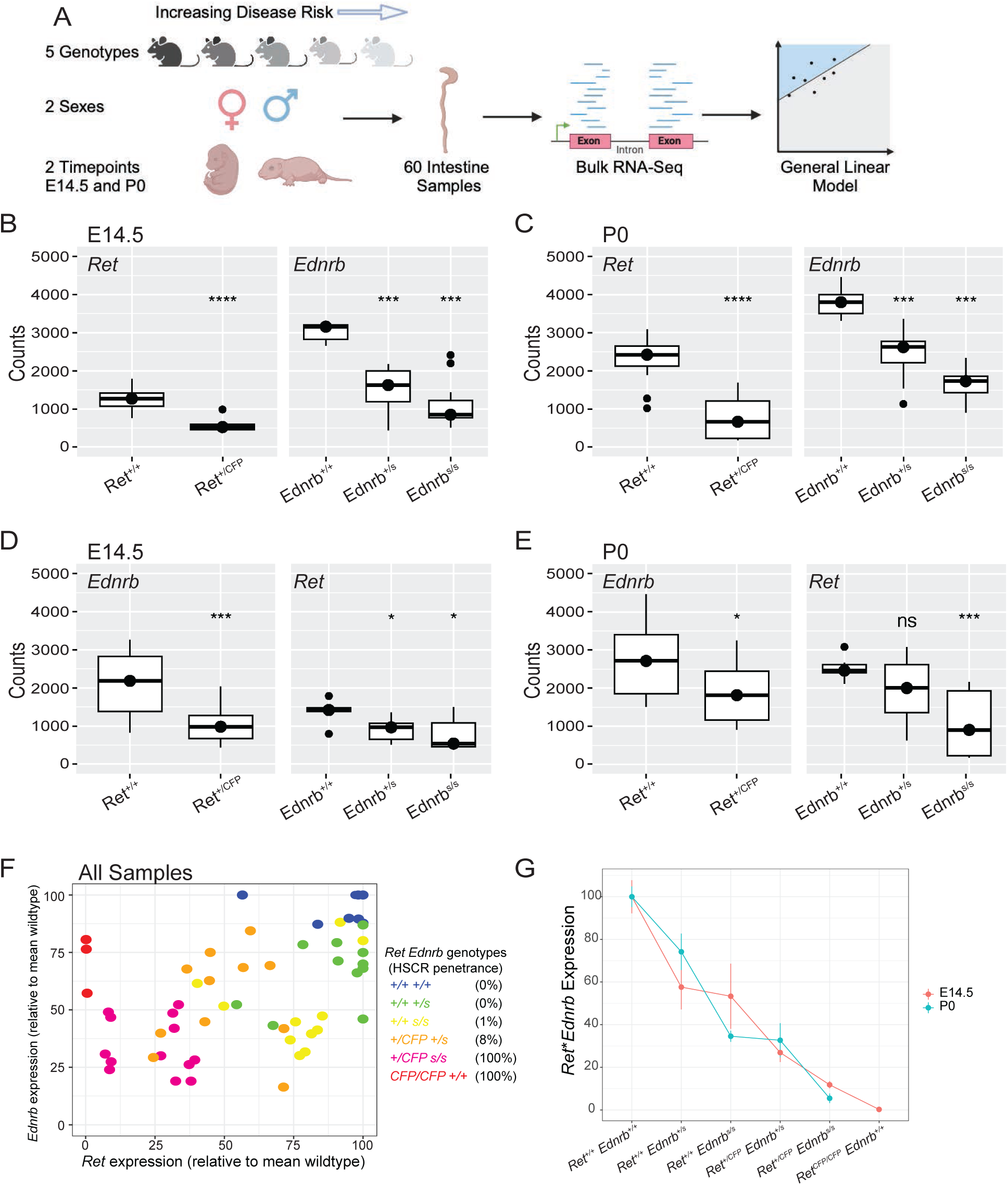
Bulk RNA-seq analysis of gut samples. **(A)** Schematic overview and statistical analysis of RNA-seq samples. Mean normalized *Ret* and *Ednrb* gene expression (read counts) of 28 E14.5 samples **(B)** and 30 P0 samples **(C)** showing linear decrease with increasing mutant allele dosage of the respective gene. Mean normalized *Ret* and *Ednrb* gene expression (read counts) of 28 E14.5 samples **(D)** and 30 P0 samples **(E)** showing linear decrease with increasing mutant allele dosage of the other gene (trans effect). **(F)** Pairwise plot of *Ret* and *Ednrb* normalized read counts of all 58 samples, together with our previously published data on 6 *Ret* null homozygotes,^18^ relative to the mean gene expression of WT samples. **(G)** Plot of the product of *Ret* and *Ednrb* gene expression by genotype and developmental time relative to the mean gene expression of WT samples.

We first performed principal component analysis (PCA) on the 58 samples and noted that developmental time accounted for 79% of the variance (**Figure S2A**). Since gene counts were on average higher at E14.5 as compared to P0, and were significantly different, we analyzed these two time points separately. This is appropriate because the effects at P0 are both from effects of *Ret* and *Ednrb* deficiency at P0 as well as effects on gut cell composition from perturbed development at E14.5. This was not the case for *Ret* dosage, *Ednrb* dosage, or sex (**Figure S2 B-E**). Upon splitting E14.5 and P0 samples, we no longer observed excess variation by PCA or of Cook’s distance between sample replicates (**Figure S3 A-C**). To further reduce noise, we filtered our gene list by excluding genes with <5 counts in any sample unless that gene had >300 counts across all samples; note that some null or small counts could be due to loss of *Ret* or *Ednrb* gene expression. We also removed 2 samples with inadequate sequencing coverage (**Figure S3D, E**). This left us with 58 samples and 12,512 genes (from an initial annotated list of 21,656 transcripts) for analysis, including 27 of 28 genes linked to HSCR from past studies.^9^

We examined gene expression of *Ret* and *Ednrb* as positive controls relative to the doses of *Ret^CFP^* and *Ednrb^s^* genotypes. As expected, individually, *Ret* and *Ednrb* normalized read counts decreased linearly with respect to *Ret^CFP^* and *Ednrb^s^* mutant allele dosage, respectively, regardless of the genotype of the other gene, sex, or timepoint (**Figure 2B, C**). Additionally, we also analyzed *Ret* (and *Ednrb*) gene expression with respect to *Ednrb* (and *Ret*) mutant allele dosage: consistent with their epistasis, each gene declined in expression with increasing mutant gene dosage of the other (**Figure 2D, E**).

The above results suggested that we study gene expression effects across the transcriptome with respect to the joint mutant allele dosages at *Ret* and *Ednrb*. Accordingly, we plotted the gene expression of each individual sample per genotype together with six *Ret^CFP/CFP^* null homozygote samples from one of our previous studies as a positive control (**Figure 2F**).^18^ Note that the five genotypes we study here can be arranged as an allelic series as *Ret^+/+^Ednrb^+/+^*, *Ret^+/+^Ednrb^+/s^*, *Ret^+/+^Ednrb^s/s^*, *Ret^+/CFP^Ednrb^+/s^*, and *Ret^+/CFP^Ednrb^s/s^*, with a combined mutant allele dosage of 0, 1, 2, 2, and 3, and disease penetrance of 0%, 0%, 1%, 8% and 100%.^19,15^ *Ret^CFP/CFP^ Ednrb^+/+^* has 100% penetrance with a mutant allele dosage of 2 although this is not true for the *Ret^+/+^Ednrb^s/s^* genotype. The general trend towards higher penetrance with decreasing expression of *Ret* and *Ednrb* is clear although individual samples demonstrate considerable variance with respect to their average levels (**Figure 2F)**. This variation may be due to either measurement variation or variation in the numbers of ENCDCs across individuals of any one genotype or both.^8^ Furthermore, the *Ednrb^s^* allele is somewhat more variable than the *Ret^CFP^* null allele, likely due to the stochastic nature of the retrotransposon mis-splicing event. Our results demonstrate that although individual animals can develop aganglionosis rarely with 2 or fewer combined mutant alleles, full penetrance requires greater mutant allele dosage at both genes. We note that the subtle loss of ENS density we observe in the cecum and colon starts in the *Ret^+/CFP^ Ednrb^+/s^* genotype at ∼30% gene expression of the product of *Ret* and *Ednrb* relative to mean *Ret^+/+^Ednrb^+/+^* expression (**Figure 2G**). Further reduction to around 10% expression of the product relative to *Ret^+/+^Ednrb^+/+^* is the threshold where 100% penetrance of long segment aganglionosis occurs, as observed at both E14.5 and P0 in the *Ret^+/CFP^ Ednrb^s/s^* genotypes (**Figure 2G**). This suggests that reduction of ENS innervation linearly correlates with reduced *Ret* and *Ednrb* expression. Additionally, our data also reveals that female *Ret* null mice had higher expression levels of *Ednrb* than male *Ret* null mice (**Figure 2F**). We did not see this trend for other genotypes, but this observation is intriguing as it is consistent with the lower disease penetrance in females than male.

Finally, we examined the expression of all 24 known HSCR genes with respect to the combined mutant allele dosage series with increasing penetrance.^9^ None of the genes demonstrate any specific relationship with the exception of *L1cam* that exhibits continual linear loss of its gene expression with respect to *Ret* and *Ednrb* loss at both E14.5 and P0 (**Figure S4**). *Ret* and *Ednrb* affecting expression of *L1cam*, which is located on the X chromosome, may be an explanation of both the increasing penetrance with increasing combined mutant allele dosage and the higher HSCR penetrance observed in males.

### Global effects of *Ret* and *Ednrb* deficiency, individually and synergistically

We next inquired which specific genes, across the transcriptome, are affected by the actions of *Ret* and *Ednrb*. To answer this question, we performed DESeq2 analysis using an additive general linear model accounting for *Ret* and *Ednrb* gene dosage and sex, at E14.5 and P0 separately (see **Methods)**. First, there are thousands of genes modified by *Ret* and *Ednrb* individually. GO analysis of these genes identifies broad biological processes with many genes previously reported in our past study on homozygous *Ret* null mice.^18^ At E14.5, *Ret* affects nearly twice as many genes as *Ednrb* alone (**Figure S5A**): *Ret*- and *Ednrb*-dependent changes are mostly associated with metabolism and growth, and protein binding and mRNA binding genes, respectively (**Figure S5C**). The ratio of *Ret* to *Ednrb* affected genes becomes larger at P0 with *Ret* affecting 3,893 genes relative to 265 by *Ednrb* alone (**Figure S5B**). Once again, metabolism genes are overwhelmingly enriched in *Ret*-dependent pathways whereas *Ednrb* affects 13 SNARE pathway and 25 axonal genes (**Figure S5C**). All genes affected by *Ret* and *Ednrb* additive action are reported in **Table S3**. The data also show that *Ednrb* has a stronger effect on gene expression as compared to *Ret*, consistent with their relative genetic risk in patients.^9^

To understand the joint effects of *Ret* and *Ednrb* we next examined those genes whose expression was altered only by their non-additive interaction. We identified 225 genes at E14.5 and 367 genes at P0 with this property (**Table 2**). Interestingly at E14.5, these genes have an expression pattern that is not individualistic but follow a specific increasing or decreasing pattern in relation to the combined mutant allele dosage increases or decreases, except in the most severe *Ret^+/CFP^ Ednrb^s/s^* phenotype (**Figures 3A, B**). Interaction-positive genes, defined with respect to the sign of the interaction regression coefficient (β_Ret*Ednrb_>0), generally have increased expression with increasing levels of *Ret* and *Ednrb*, while interaction-negative genes (β_Ret*Ednrb_<0) are the opposite. We hypothesize that these interactions between *Ret* and *Ednrb* are genes affecting the cellular properties of ENCDCs, and, thereby, ENS development. The likely cellular phenotype affected is proliferation and migration.^7,13^

**Figure 3.**
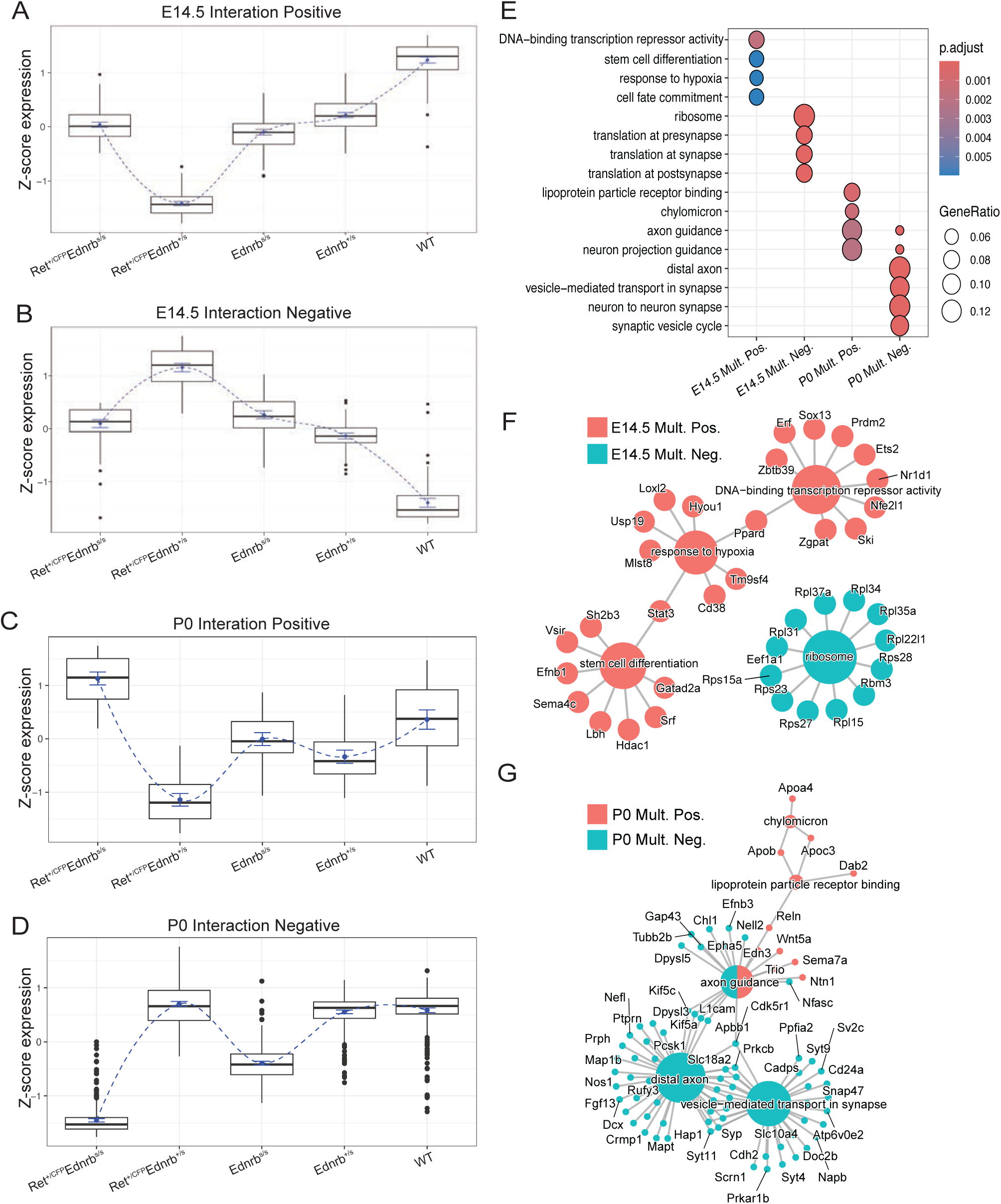
Transcriptional consequences of *Ret* and *Ednrb* interactions. Z-score boxplots at E14.5 **(A, B)** and P0 **(C, D)** of normalized counts of genes with positive and negative non-additive interactions across *Ret* and *Ednrb* genotypes. **(E)** GO analysis of genes with statistically significant (P_adj_ < 0.05) gene expression alterations from *Ret* and *Ednrb* interactions using the clusterProfiler package; only the top 4 terms are plotted for each significant genes class (X-axis). Cnetplot indicating the major genes at E14.5 **(F)** and P0 **(G)** driving the GO annotation terms.

**Table 2.** Significant Genes. Total number of significantly affected genes (p<.05) by either *Ret* alone, *Ednrb* alone, their additive effect, or multiplicative interaction from our linear regression model (see Methods).

| Coefficient | E14.5 | P0 |
| --- | --- | --- |
| *Ret* Pos. | 1 | 18 |
| *Ret* Neg. | 0 | 16 |
| *Ednrb* Pos. | 0 | 54 |
| *Ednrb* Neg. | 0 | 8 |
| *Ret* + *Ednrb* Pos. | 0 | 13 |
| *Ret* + *Ednrb* Neg. | 0 | 2 |
| *Ret* * *Ednrb* Pos. | 135 | 52 |
| *Ret* * *Ednrb* Neg. | 90 | 315 |

Through annotation analysis, we note that there is an enrichment of chromatin modifying genes (e.g., *Hdac1*) in the positive interaction gene set as well as several stem cell differentiation genes (**Figure 3E**). In particular, *Hdac1* has been recently suggested to be at the core of a genetic response to HSCR in a scRNA-seq study of human patient-derived cell lines (**Figure 3F).**^23^ While we see a similar trend for P0 interaction positive genes, it is noteworthy that the relationship between genotype and gene expression is much more muted at P0 than at E14.5 (**Figures 3C, D**). We assume that this effect is owing to the consequences of aganglionosis itself rather than its specific molecular etiologies. This is most evident by the P0 interaction negative genes showing extensive enrichment (67/315 or 21%) for neuronal genes (**Figures 3D, G**). At P0 there are several genes still affected by *Ret* and *Ednrb* individually in our analysis. This is most likely due to the separation of *Ret* and *Ednrb* expression into distinct lineages beyond E14.5 when ENS development is normally completed.^22^

### Non-cell autonomous effects of *Ret* and *Ednrb* interaction

Given the above observations, we wanted to assess whether the genes specifically altered by *Ret* and *Ednrb* interaction were resident in the same cells or not, i.e., were they cell autonomous or not? However, given the general sparsity of genes with detected expression in single cells we paired this analysis with our prior bulk RNA-seq data.^24,25^ We isolated cells from E14.5 *Ret^+/+^Ednrb^+/+^* embryonic tissue from the small intestine to the colon, filtered the data (**Methods**), and annotated cell clusters according to the literature as well as our own studies (**Figures 4A-C**).^26–29^ Next, we assessed whether there was any enrichment for these gene classes within specific populations of cells (**Figures 4D, E**). We estimated the module score which measures the expression enrichment of a predefined group of genes in comparison to 100 randomly chosen control genes from the same cluster of cells per gene in the predefined group: a negative or near zero score indicates non-enrichment or low expression in that cell type. Interestingly, despite increasing gene expression in the *Ret^+/CFP^ Ednrb^s/s^* mutant, positive *Ret* and *Ednrb* interacting genes are depleted in neurons whereas negative interaction genes are enriched in neurons, glia, and ENS progenitors (**Figures 4D, E**). This is somewhat paradoxical because if we assume that the enteric nervous system tissue is diminished in the *Ret^+/CFP^ Ednrb^s/s^* mutant, then we would predict the opposite.

**Figure 4.**
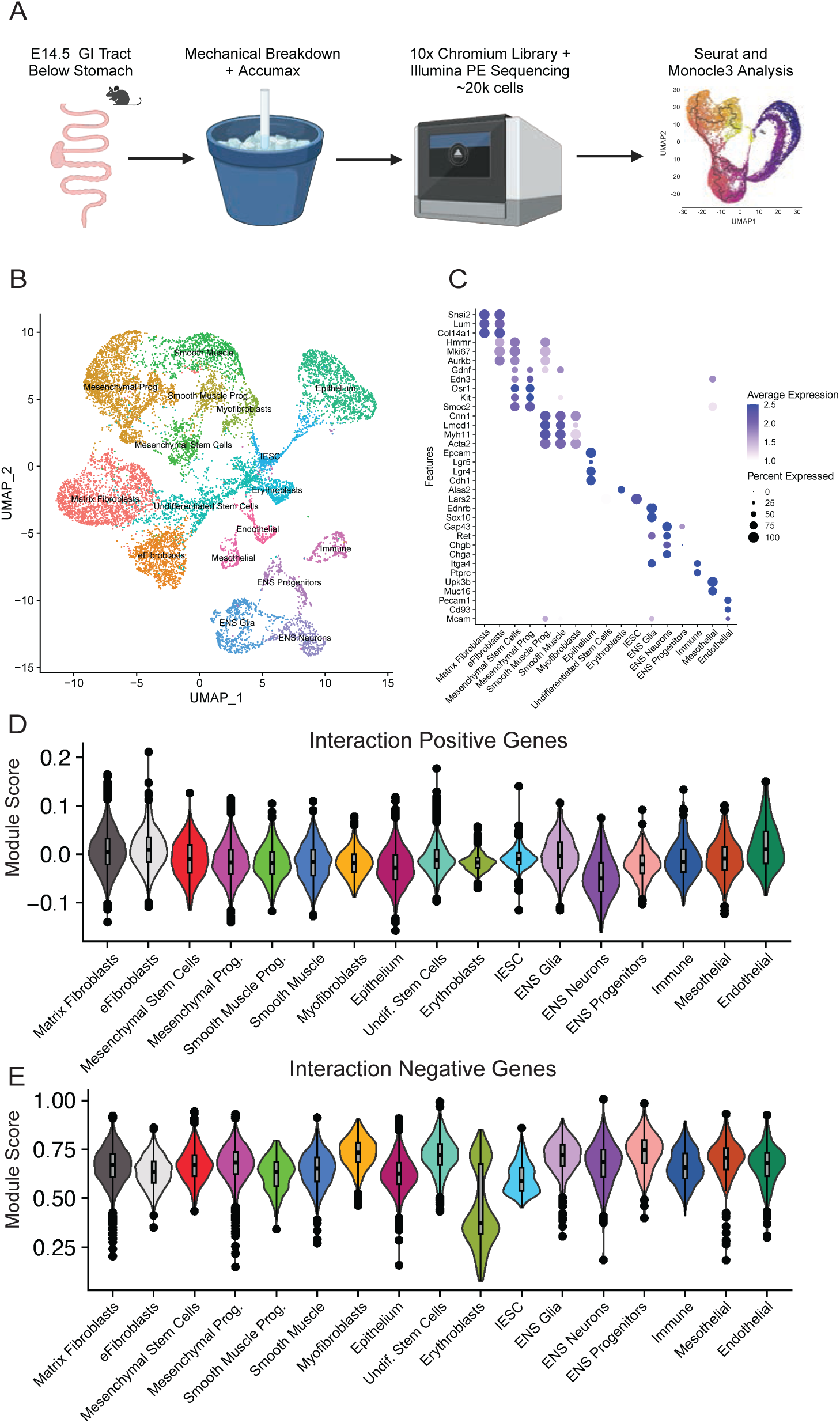
Single cell gene set analyses from gene expression data. **(A)** Overview of single cell isolation, library generation, and scRNA-seq analysis. **(B)** UMAP display of WT mouse gut samples. **(C)** Dotplot of the average gene expression of markers used to identify cell types. **(D, E)** Violin plots of gene expression module scores for the positive and negative interaction gene sets at E14.5 for 17 pre-defined cell types in the WT.

To resolve this issue, we performed scRNA-seq on the piebald (*Ret^+/+^Ednrb^s/s^*) and *Ret*-deficient piebald (*Ret^+/CFP^Ednrb^s/s^*) mouse models at E14.5, in duplicate samples, to assess changes in cell populations across genotypes. We discovered that the *Ret^+/CFP^ Ednrb^s/s^* genotype had a significant reduction in neurons/glia as compared to WT, but there was an increase in the number of ENS progenitors in *Ret^+/CFP^ Ednrb^s/s^* embryos as well (**Figure 5A**). We overlayed a subset of cells of this data with *Fabp7* and *Vamp2* as representative genes for our interaction negative and interaction positive gene sets, respectively (**Figures 5C, D**). *Fabp7* expression is restricted to ENS progenitors in the *Ret^+/+^ Ednrb^+/+^* genotype (**Figure 5C, left panel**), but in the *Ret^+/+^ Ednrb^s/s^* mutants some mesenchymal cells begin to express *Fabp7* (**Figure 5C, middle panel, gray arrows**). This pattern ceases to exist in the high penetrance *Ret^+/CFP^ Ednrb^s/s^* mutant (**Figure 5C, right panel**), thereby showing greater resemblance to the WT pattern (**Figure 5C, left panel**). This is precisely what we observed in the bulk-RNA data (**Figure 3B**). There also appears to be a significant increase in *Vamp2* expression in mesenchymal cells (**Figure 5D, right panel, gray arrow**) in the *Ret^+/CFP^ Ednrb^s/s^* mutant, which could also explain the increased expression observed for this genotype in the bulk data (**Figure 3A**). To quantify these results, we pseudo-bulked the gene expression of each genotype for this subset of cells, with resultant patterns that precisely matched those in our bulk-RNA-seq dataset (**Figure 5F**).

**Figure 5.**
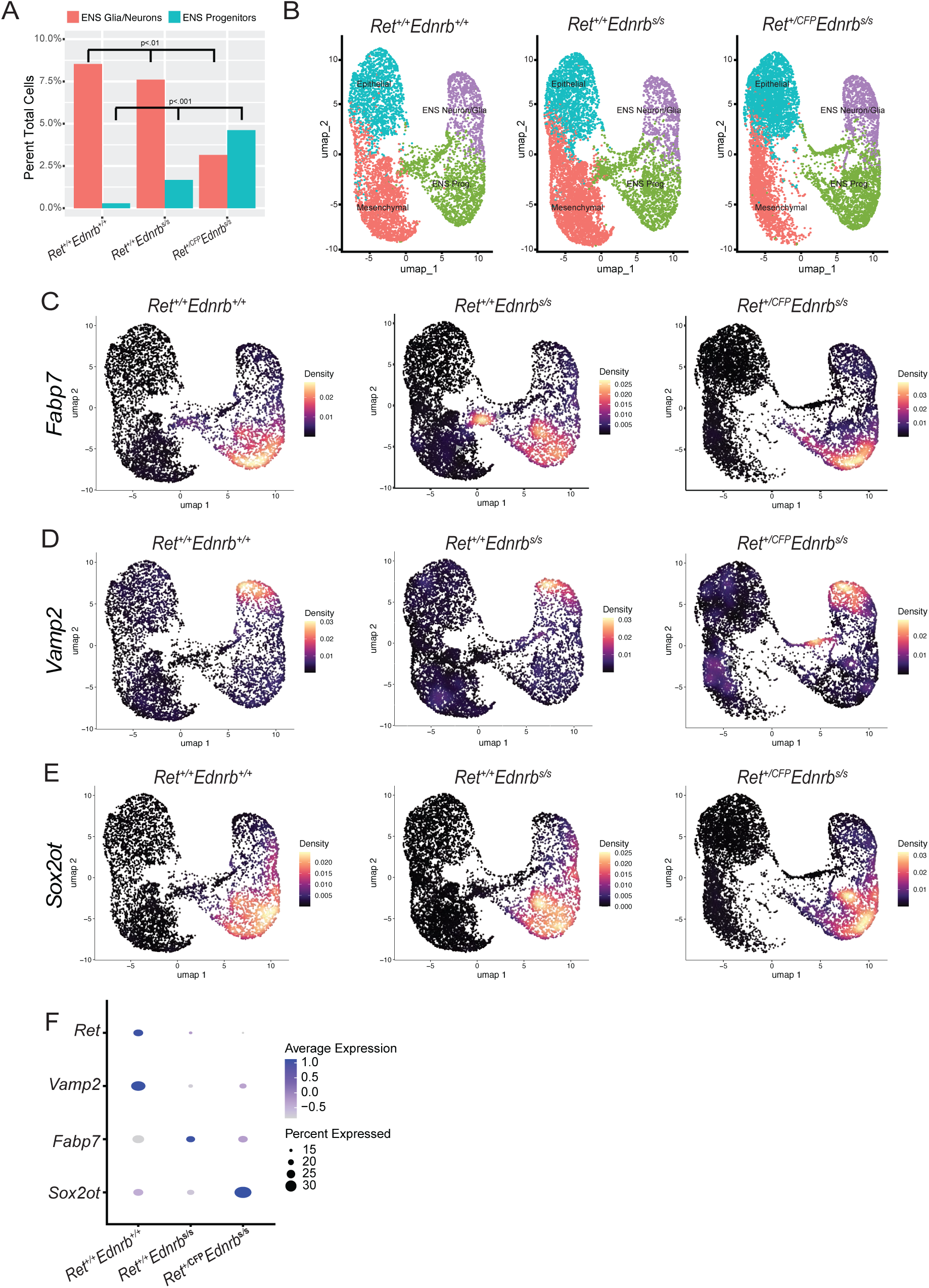
RNA-seq interaction genes reveal cell fate changes within a perturbed ENS. **(A)** Quantitation of maturing enteric neurons and glia vs. progenitor cell populations based on Seurat scRNA-seq clustering. **(B)** UMAP display of cells from all three *Ret-Ednrb* mutant genotypes with aganglionosis. **(C, D)** Density plots of *Fabp7* (representative interaction negative gene) and *Vamp2* (representative interaction positive gene) gene expression across 3 genotypes from least to most penetrant. **(E)** Density plot of *Sox2ot* gene expression across 3 genotypes from least to most penetrant. **(F)** Dotplot of pseudo-bulk data from the 3 genotypes for selected representative genes.

These analyses suggest that the expression patterns of both sets of interaction gene sets can be explained by their ectopic expression in mesenchymal cells in response to ENS developmental perturbations. We further hypothesize this may be an epithelial to mesenchymal transition (EMT) in cell fate and perhaps a programmed response for maintaining normal ENS development in response to genetic perturbation. This hypothesis is supported by an apparent increase in the number of intermediate “mesenchymal” cells in the *Ret^+/+^ Ednrb^s/s^* mutant (**FigureS6A**) as well as a decrease in the pseudo-time distance between epithelial and mesenchymal cells in *Ret^+/+^ Ednrb^s/s^* mutants versus WT mice (**Figure S6B, C**). We interpret this as a ‘rescue’ mechanism owing to the non-linear nature of our interaction genes, i.e. our interaction gene sets do not continually increase or decrease across all genotypes as would have been expected from mutant gene dosage alone. In other words, we would have expected the epithelium and mesenchyme to continually decrease gene expression in a linear manner if development of all 3 tissues was stalled with increasing ENS perturbation. This potential rescue mechanism presumably breaks down in the *Ret^+/CFP^ Ednrb^s/s^* mutant due to the severe loss of ENS development in the colon, signaling required to drive transition of other tissues in the less severe models.

Finally, we found that one of the most differentially expressed genes between our genotypes was *Sox2ot* which exhibited the highest expression in *Ret^+/CFP^ Ednrb^s/s^* mutants (**Figure 5E, F**). *Sox2ot* has been previously shown to repress neural progenitor proliferation, promote differentiation, and negatively regulate the renewal of stem cells.^30^ All of these phenotypes match what we see in the mouse, clearly suggesting that loss of *Ret* and *Ednrb* expression prevents ENS precursor cells from maintaining their ‘stemness’. So, although there is an apparent increase of progenitors in the *Ret^+/CFP^ Ednrb^s/s^* mutant (**Figure 5A**), they are differentiating far too early as compared to the same cell populations in other genotypes.

## Discussion

### Null mouse models are inadequate for modeling a complex disorder

In this study, we hypothesized that mouse models with jointly decreasing levels of *Ret* and *Ednrb* gene expression in the developing gut would be necessary to gain novel insights into HSCR disease mechanisms, in contrast to complete knockout gene models.^18,19^ Indeed, we observed several human-like phenotypes in our mouse models: the presence of megacolon at P21; subtle and variable enteric neuron density defects in the cecum of otherwise ganglionic compound heterozygous *Ret^+/CFP^ Ednrb^+/s^* mice; and, higher penetrance in males. Our gene expression analysis of the developing gut also revealed an extensive upregulation of metabolic genes involved in mitochondrial and ribosome function at both E14.5 and P0, a feature not revealed in our prior studies of *Ret* homozygous null mice.^18^ Although both studies identified HSCR candidate genes, the classical null models were insufficient for deciphering the entire gene universe and cellular outcomes relevant to phenotypes of human patients. Our data also reveals the most likely reason for *Ednrb* affected gene enrichment in neuronal terms and higher genetic risk as compared to *Ret* in human genetic studies.^9^ Namely, neurons are highly sensitive to *Ednrb* dosage due to the already naturally low levels of *Ednrb* levels in WT neurons (**Figure 4C**), with a greater impact from *Ednrb* risk variants.

The data from this study also offer a unique resource for querying candidate genes from patient cohorts. Indeed, from our recent rare variant case-control exome analysis of 301 probands, we identified 2 patients with pathogenic variants in *TMEFF2*, which is not a statistically significant excess over controls.^36^ Yet, *Tmeff2* is differentially affected by *Ret* and *Ednrb* in our bulk-RNA data and exhibits enriched expression in enteric neurons in our single cell data implicating the gene in HSCR.^31^ These data also raises the intriguing possibility that one cause of the male excess in HSCR may be from *Ret* and *Ednrb*’s interaction effect on *L1cam.* This gene, located on the long arm of the chromosome X, is the only sex-linked gene with decreased expression with increasing genetic risk across all of our genotypes and at both timepoints (**Figure S4**). *L1cam* is a known HSCR gene,^3^ and could have analogous effects without coding variants if *Ret* and *Ednrb* were to reduce its gene expression and be a cause of its male sex bias.

### Gene Regulatory Networks and rate-limiting rheostats

We and others have postulated that GRNs are the simplest units of cell function and disease, with good evidence for this in HSCR. Beyond obvious biochemical interactions, GRNs allow for reducing transcriptional noise arising from fluctuation in transcripts and protein levels of GRN member genes that may be due to genetic variation, epigenetic fluctuations, chromatin remodeling, environmental changes, or cell cycle state over time, particularly during the rapid changes occurring in development.^32^ GRNs can also account for disease penetrance differences when one considers deleterious mutations at core rate-limiting genes vs external nodes. In this sense, *Ret* and *Ednrb* appear to be rate-limiting genes that link an extensive number of upstream ligands and transcription factors with downstream signaling pathways in ENCDCs. However, are *Ret* and *Ednrb* transcript levels, and other members of the core GRN, sufficient to explain HSCR phenotypes or do we require knowledge of the entire HSCR gene universe? In other words, can there be a second core GRN or other sets of variants outside of the *Ret-Ednrb* GRN or downstream of their signaling partners that can produce HSCR without perturbing *Ret* or *Ednrb* expression? The current data suggests not. Future work should attempt to discern if HSCR can be defined simply by expression of *Ret* and *Ednrb* as this may prove useful in diagnosis.

### Robust cell fate changes in response to developmental insult

HSCR has long been phenotypically characterized as the absence of neuronal ganglion cells. However, 80% of patients are affected only in the very distal part of the colon with so called short-segment HSCR.^33^ In other words, the entire small intestine and majority of the colon in most probands is phenotypically normal. Our bulk and single cell RNA analyses suggest that there is an alteration in the developmental fate of cells in the developing gut when genetically perturbed by reducing *Ret* and *Ednrb* expression. Therefore, HSCR is not solely a loss of enteric neurons. This opens the possibility that there are additional pathways and variants that can increase or decrease the burden threshold of deleterious variants in *Ret* or *Ednrb* in a non-cell autonomous manner. Such an intricate relationship between the ENS, mesenchyme, and epithelium has been previously characterized. In zebrafish, there are epithelial mutants that can affect both smooth muscle and ENS development.^34^ In chicken embryos, inhibition of smooth muscle development can also drastically reduce ENS radial innervation of the gut without affecting proliferation or increasing apoptosis.^35^ Conversely, ENS ablation before GI tract colonization prevents smooth muscle development of the stomach in chicken embryos.^36^ Our data suggest that a subset of immature cell populations of the mesenchyme and epithelium may shift towards a neuronal fate in order to support initial perturbations to ENS development (**Figures 5, S6**). Future effort should be geared to determine if this is the case or if it is a non-cell autonomous effect where the entire GI tract fails to develop normally due to lack of proper ENS developmental cues. This alternative hypothesis would explain why epithelial cells are closer in pseudotime to mesenchymal and neuronal progenitor cells in the *Ret^+/+^ Ednrb^s/s^* mutant versus WT **(Figures S6B, C)**, but it doesn’t explain the non-linear relationship between our interaction genes and the perturbation of *Ret* and *Ednrb* **(Figures 3A, B)**.

### Materials and Methods Mouse strains used

The *Ret-CFP* knock-in mouse line, in which exon 1 of *Ret* has been replaced by a cyan fluorescent protein (CFP) cDNA sequence followed by a polyA tail, was previously made and extensively characterized, by others and us, in earlier work.^13 18,16^ The *Ednrb-s* (*piebald*) mouse line (stock no. 000179-MU)^37^ was obtained from frozen sperm through the Mutant Mouse Resource & Research Center (MMRRC) consortium. All procedures were approved by the New York University Animal Care and Use Committee (protocol number: IA17-01779). Mice of both sexes were used unless specified otherwise.

### Mouse breeding and genotyping

Homozygous *Ret^CFP/CFP^* mice are lethal within 1 week of birth due to kidney defects.^13^ Thus, heterozygous *Ret^CFP/+^* mice were crossed with homozygous *piebald Ednrb^s/s^* mice to produce F1 compound heterozygote *Ret^CFP/+^, Ednrb^s/+^* mice. These compound heterozygotes were next crossed to homozygous *piebald Ednrb^s/s^* mice to generate four of the *Ret-Ednrb* mouse genotypes studied here. DNA from adult mice was extracted from an ear-punch biopsy, while DNA from newborn (P0) and embryonic mice were obtained from a tail snip and forelimb, respectively. All tissue was broken down in Proteinase K digestion buffer (50mM Tris pH7.5, 5mM EDTA, 1% SDS, 0.2M NaCl, 0.2mg/mL Proteinase K) for 2-3 hours at 55°C followed by a 1:1 phenol: chloroform organic-phase clean-up and ethanol precipitation of DNA. Routine genotyping was performed by PCR using the following primers: universal reverse primer 5’-CAGCTAGCCGCAGCGACCCGGTTC-3’, with either forward wildtype *Ret* primer 5’-CAGCGCAGGTCTCTCATCAGTACCGCA-3’ resulting in a 550bp sized band, or forward *Ret-CFP* detection primer 5’-ATCACATGGTCCTGCTGGAG-3’ resulting in a 449bp sized band. The *piebald* allele was genotyped in a similar manner with universal reverse primer 5’-GAGTTTGCTTCCAACCAGTG-3’, with either forward wildtype *Ednrb* primer 5’-CAAGAAGGTAGTATAGCCAGGAG-3’ resulting in a 550bp sized band, or forward *piebald* detection primer 5’-ACTCCCATTTCCTTCCAGGT-3’ resulting in a 500bp sized band. Sex was assessed using *Jarid* forward 5’-CTGAAGCTTTTGGCTTTGAG-3’ and reverse 5’-CCGCTGCCAAATTCTTTGG-3’ primers from previously published work,^38^ resulting in a single 331bp band in XX females or an additional band of 301bp in XY males.

### Acetylcholinesterase staining for phenotyping

P0 mice were euthanized with isoflurane in a closed chamber followed by decapitation to ensure death. The entirety of the gut, from duodenum to the anal pore, was dissected out of euthanized P0 mice and immediately placed into cold 1X Phosphate buffered saline (PBS). A Stemi SV6 stereo dissection microscope was used to carefully examine the tissue and remove excess mesentery and fatty tissue surrounding the intestine. To improve image quality, select samples where gently massaged with fine tweezers to remove fecal matter. 16% methanol-free formaldehyde ampules were diluted to 4% in PBS and used to fix tissue for 1-2 hours at room temperature (RT) on a Nutator. Tissue was then placed in a 1.72M saturated sodium sulfate solution overnight at 4°C. The following day, the tissue was incubated for 4 hours in substrate buffer (0.2mM Ethopropazine HCl, 4mM Acetylthiolcholine iodide, 10mM Glycine, 2mM Cupric sulfate pentahydrate, 65mM Sodium acetate) at RT. Finally, a small aliquot of 1.25% sodium sulfide pH6.0 was added to develop the stain at RT for 5 minutes. The tissue was washed with dH_2_O several times and photographs were taken with a trans-illumination on a stereomicroscope using whole mounts in water.

### Fluorescence (CFP) imaging

P0 mice and intestinal tissue was dissected and immediately analyzed using a Zeiss Axiozoom V16 fluorescence stereomicroscope with a CFP filter to take photographs of whole mount intestinal tissue in cold 1X PBS. E14.5 embryos were processed in a similar manner with individual mothers euthanized by isoflurane exposure followed by cervical dislocation. In certain cases, z-stacks were taken, and orthogonal projections were created to visualize 3D structures, most notably the cecum.

### RNA extraction and bulk RNA-seq

E14.5 (stomach to anal pore) or P0 (distal colon) gut tissue was dissected on a stereomicroscope, rinsed in cold 1X PBS, and immediately snap-frozen in liquid N_2_. For E14.5, sample tissue was mechanically lysed in TRIzol (Life Technologies) using a single 5mM stainless steel bead and a Qiagen Tissuelyser II machine for 30 secs at 20 Hz. Chloroform was added to each sample and spun down in a minicentrifuge. The aqueous phase was collected, and RNA was precipitated with isopropanol. RNA was washed in 80% ethanol and resuspended in 1X TE. DNA was removed using a DNA-free kit (Thermofisher) using the manufacturer’s instructions. For P0, the above steps were similar except a Qiagen RNeasy plus kit was used to extract RNA. Illumina Truseq cDNA libraries were prepared using an automated Beckman Coulter Biomek SPRIworks platform by the NYU Genome Technology Center. Libraries were barcoded for multiplexing and sequenced on an SP100 flow cell (2×50 paired end reads) on an Illumina Novaseq 6000.

### RNA-seq data analysis

Raw paired-end reads were processed using an automated Salmon pipeline, Seq-N-Slide,^39^ on New York University’s Ultra Violet computing cluster. Seq-N-Slide is available at https://github.com/igordot/sns. Ensembl stable IDs were then mapped to gene IDs with lncRNAs, miRNAs, pseudogenes, etc. filtered out by using the Gencode mm10 M28 annotation release to select for protein coding genes only. The final cleaned counts are available in Supplementary **Table S1** along with the sample metadata in **Table S2**. Genes with <5 counts in a replicate were filtered out unless they possessed >300 counts summed across all 58 samples. Two samples were removed during analysis due to unexpected low sequence coverage but have been submitted to GEO for potentially useful preliminary analysis. Cleaned counts were then processed through DESeq2^40^ along with two general linear models split by time (of the form):

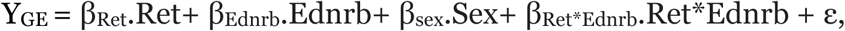

where “ is normally distributed error. For the regressors, Ret and Ednrb were given nominal values relative to their respective gene expression in our dataset (E14.5: 100% Ret-4, 50% Ret-2, 100% Ednrb-4, 59% Ednrb-2.4, 33% Ednrb-1.3) and (P0 100% Ret-4, 58% Ret-2.3, 100%-Ednrb-4, 50% Ednrb-2, 25% Ednrb-1). Sex was given a binary value of 0 (female) or 1 (male) with one model created for each time point using DESeq2 as the backbone for statistical analysis.^40^ GO and Cnet plots were produced with the clusterprofiler R package,^41^ and additional plots were produced using ggplot2^42^ with custom scripts.

### Single cell isolation and sequencing analysis

E14.5 tissue was dissected from the small intestine to the colon and rinsed in 1X Phosphate Buffer Saline (PBS) on ice. Tissue was then singularized by addition of 1mL of Accumax dissociation solution and simple pipetting/mechanical grinding through 100uμM and 40μM filters in succession. Filters were washed with an additional 1mL of 1X PBS 1% FBS. Cells were counted using a hemocytometer and diluted or concentrated such that ∼10-20,000 cells were used for standard 3’ end 10X Genomics scRNA library preparation. All single cell libraries were sequenced on an Illumina NovaSeq6000 to a minimum of ∼500m reads. For genotyping, we used embryonic tail tissue and the KAPA mouse genotyping kit (Roche) with reduced PCR cycling to genotype in under 3 hours. Data were primarily processed in Seurat,^43^ filtering for cells with at least 500 Unique Molecular Identifiers (UMIOs), at least 250 Genes, a log_10_GenesPerUMI ratio greater than 80 percent and a mitochondrial read ratio of less than 20 percent. The cell matrix was then normalized using scTransform.^44^ Violin plots of Module Scores and density plots of representative genes were produced with scCustomize.^45^ Trajectory analysis was performed with the Monocle3 package.^46^ Pseudo-counting of representative genes was done with Seurat’s AggregateExpression function on the subset of cells presented and values were plotted using Seurat’s Dotplot function.

## Data availability

All raw read data and metadata have been deposited into NCBI’s GEO database under a super series accession number GSE255964.54. Bulk RNA-seq files are under GSE227574 accompanied by Bulk-RNA-seq cleaned counts files and metadata files that have been deposited as **Table S1** and **Tables S2**. E14.5 WT and *Ret* null samples are from GEO dataset GSE103070. scRNA-seq files are deposited under the sub-series accession GSE255963 with an accompanying annotated Seurat objects in rds format.

## Acknowledgements

We thank members of the Chakravarti lab and the NYULH Center for Human Genetics and Genomics for scientific critique and discussions. Experimental schematic Figures 1A, 2A, and 4A were created in Biorender through a license to NYULH. We appreciate the efforts of the NYU Genome Technology Center for making the RNA-sequencing libraries and sequencing on the Illumina platforms. THE NYU-GTC is supported by the NYU Cancer Center Support Grant P30CA016087. This project was funded by NIH grant HD028088 and other internal funds awarded to Aravinda Chakravarti. We declare no financial or other conflicts of interest.

## Notes

### Competing Interest Statement

The authors have declared no competing interest.

