## Supplemental Figures for "Joint disruption of *Ret* and *Ednrb* transcription drives cell fate reversal in the Enteric Nervous System in Hirschsprung disease"

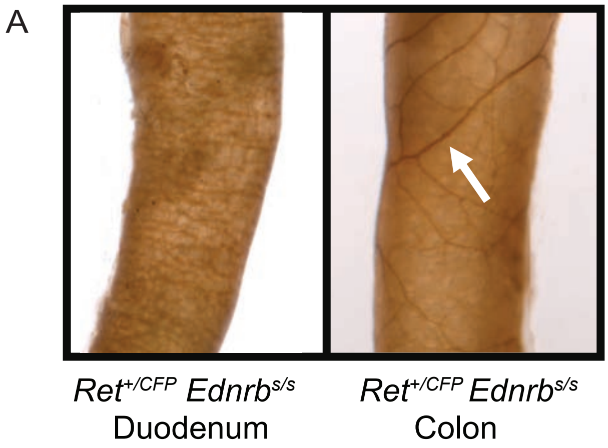


**Figure S1. The *$Ret^{+/CFP}$ $Ednrb^{s/s}$* mouse exhibits partial aganglionosis restricted to the colon (A)** Representative AChE staining of *$Ret^{+/CFP}$ $Ednrb^{s/s}$* mouse duodenum (left panel) and colon (right panel). White arrow highlights the elongated and sparse neurons in the aganglionic tissue.

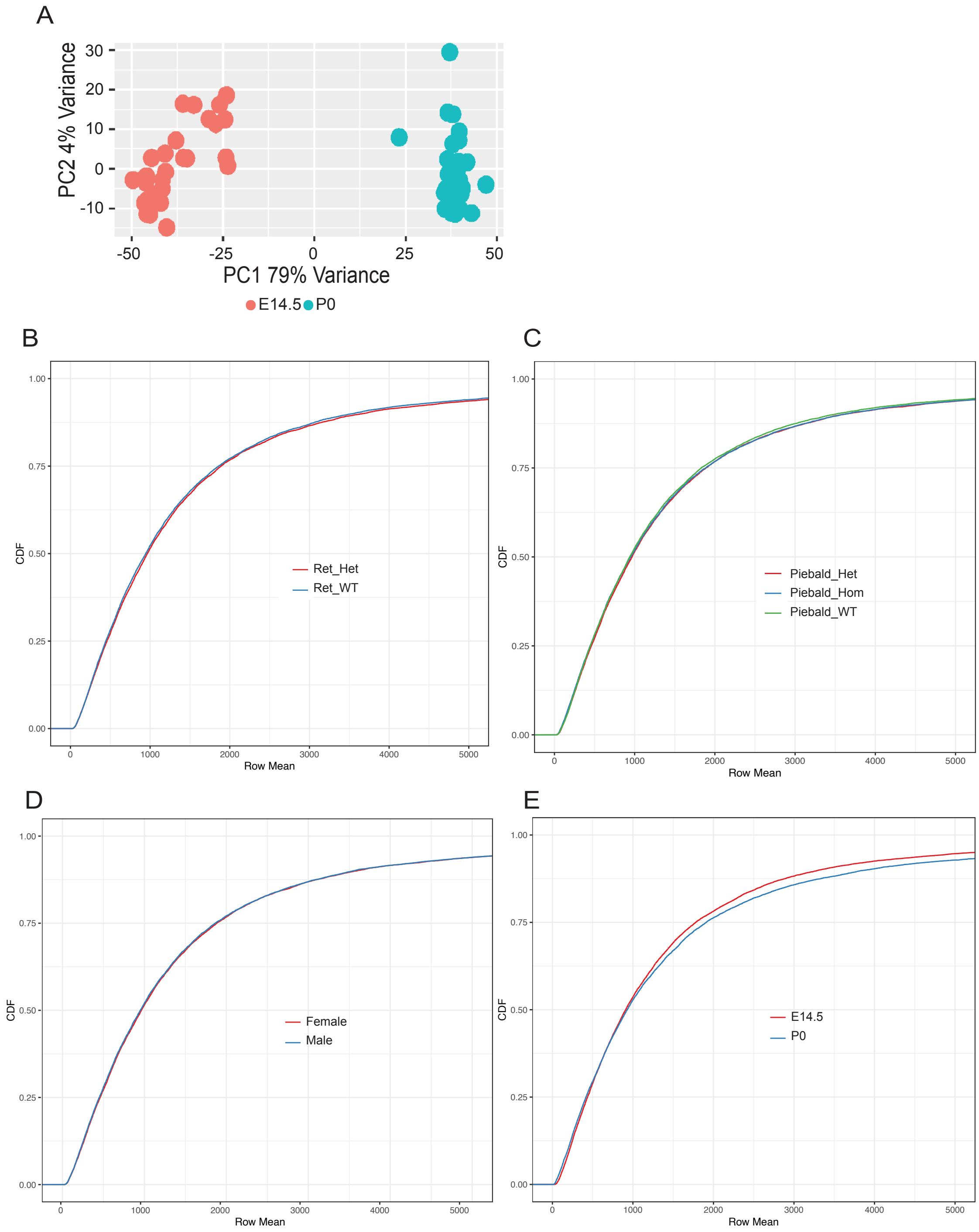


**Figure S2. Principal component analysis (PCA) and cumulative distribution functions (CDF) of mean read counts of genes by covariates reveals that time cannot be linearly modelled (A)** PCA plot showing that developmental time accounted for 79% of expression variance across time. CDF for **(B)** *Ret* genotypes, **(C)** Ednrb genotypes, **(D)** sex, and **(E)** developmental time. Only developmental time shows significant differences in the CDF with genes at E14.5 being, on average, of higher expression than at P0.

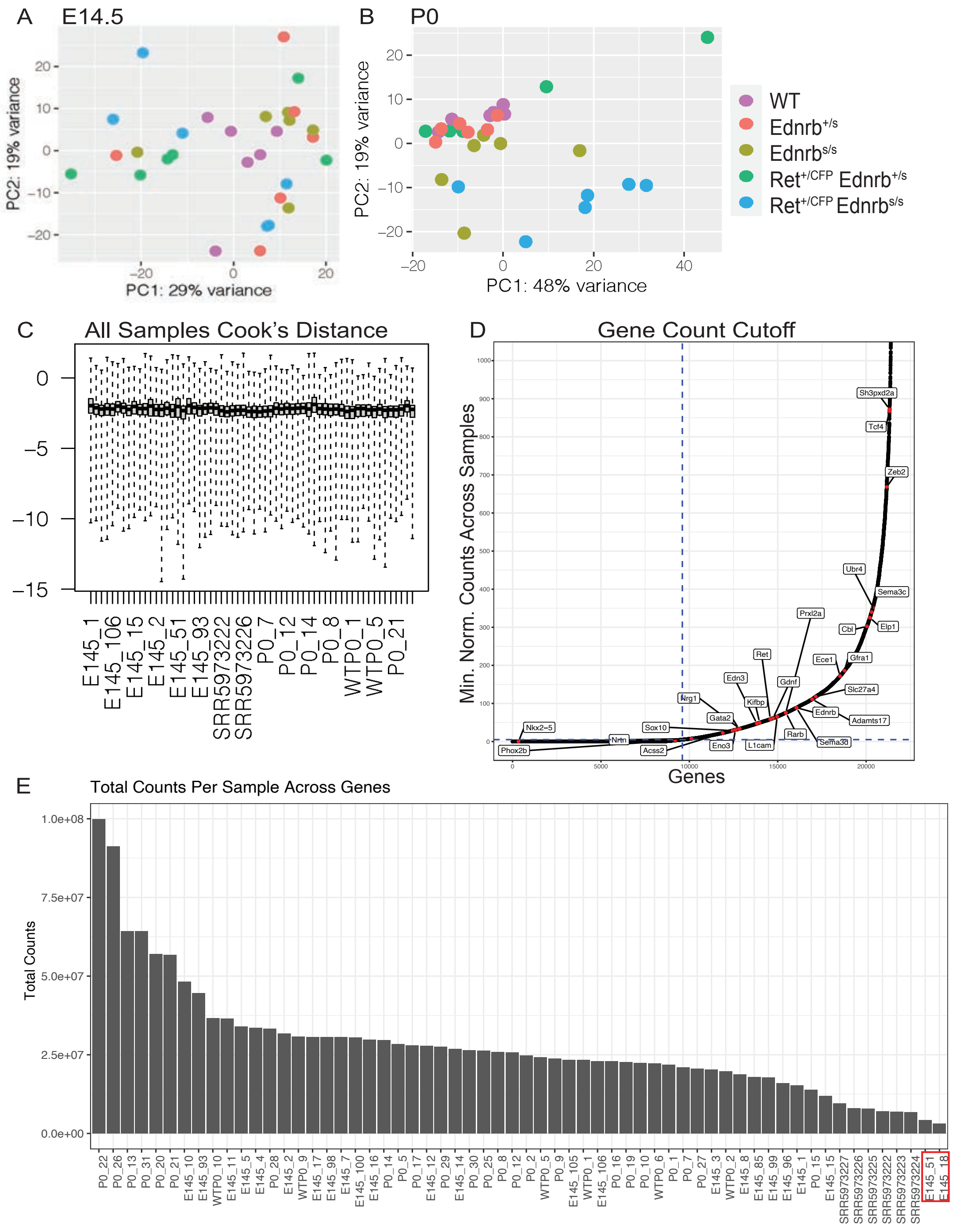


**Figure S3. Additional quality control analyses prior to regression analysis**

PCA of the **(A)** 28 E14.5 and **(B)** 30 P0 samples show no genotype associations within each developmental stage. **(C)** Box plots of Cook's distance detect no outliers within replicates or between samples. **(D)** Distribution of minimum gene expression counts for all genes across all samples was used to determine read count cutoff threshold. Genes on the bottom left have at least one sample with very few counts whereas genes on the top right have high counts in all samples. Previously reported HSCR genes from past studies are boxed. **(E)** Distribution of total read counts per sample for all genes; we excluded the two low-coverage libraries E145_51 and E145_18 from further analysis.

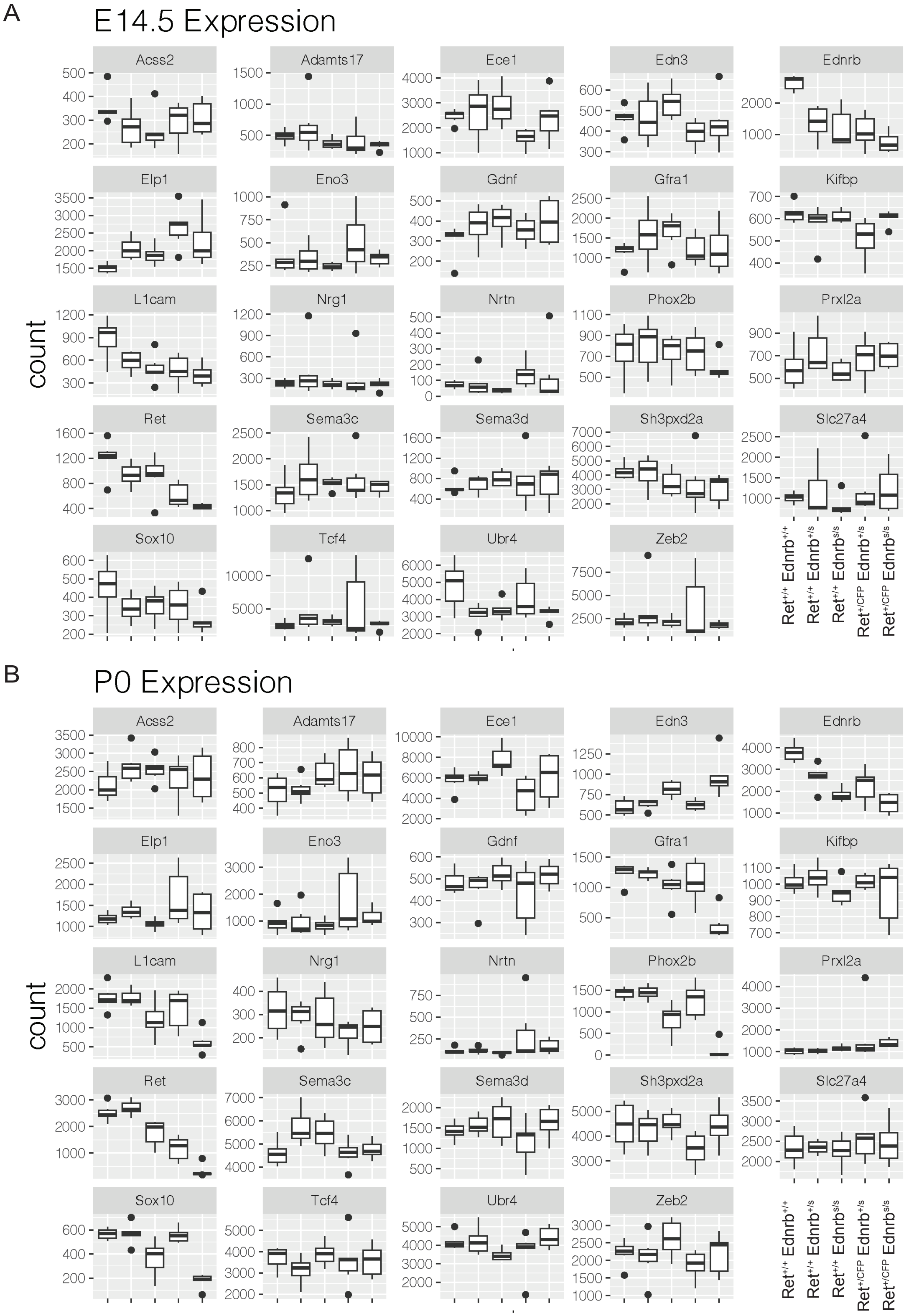


**Figure S4. Gene expression profiles of 24 HSCR genes from Tilghman et al.**[14] Boxplots of mean gene expression read counts at E14.5 **(A)** and P0 **(B)** across the 5 genotypes studied, arranged left to right from the lowest to the greatest risk of aganglionosis.

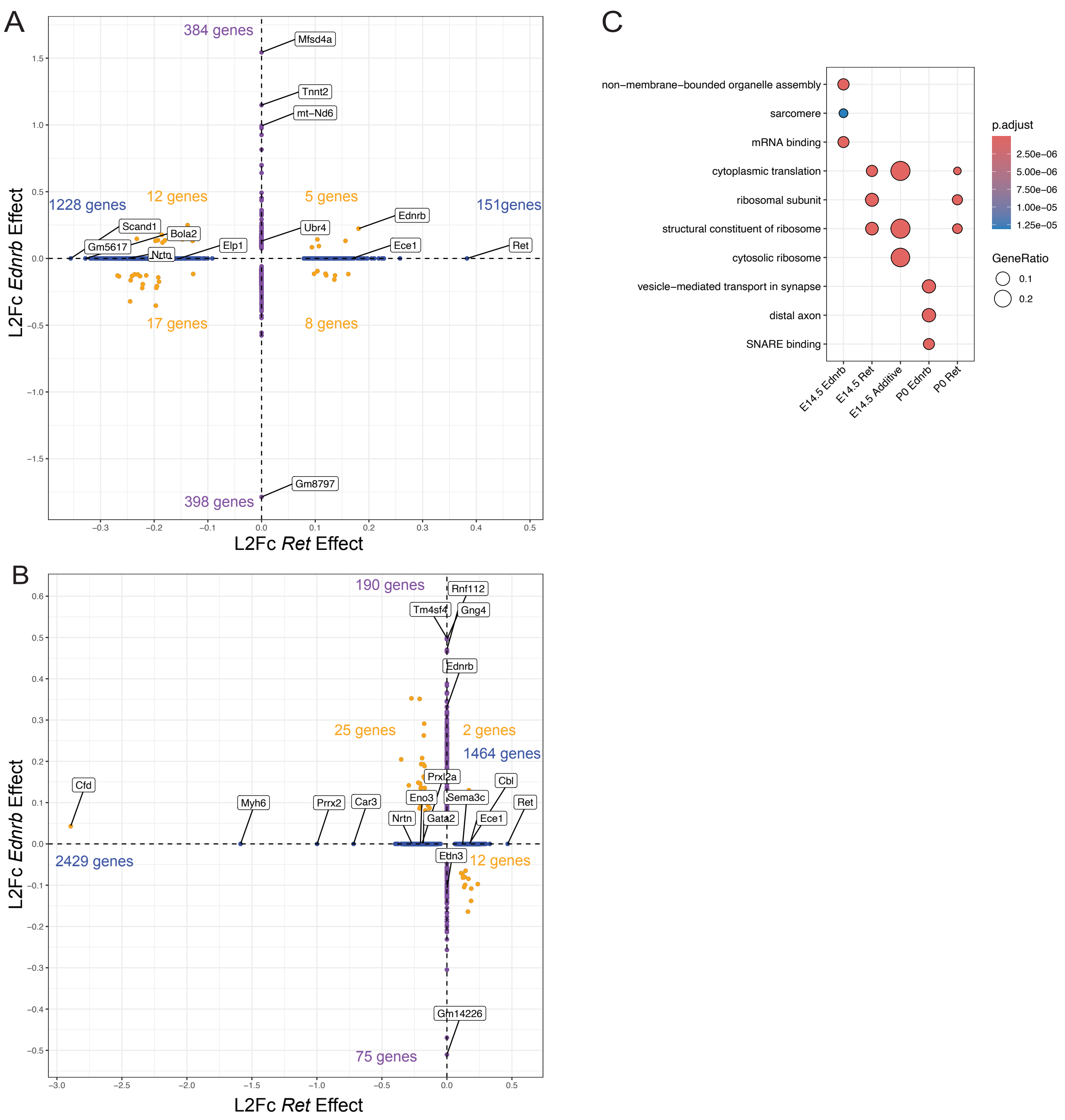


**Figure S5. Results of *Ret* and *Ednrb* additive gene expression effects**

Scatterplot of all $\log_2$-fold change values of gene expression from independent *Ret* and *Ednrb* effects at E14.5 **(A)** and P0 **(B)**. Blue-colored genes were affected by *Ret* alone, purple-colored genes by *Ednrb* alone, and orange-colored genes by both in an additive manner. **(C)** GO analysis showing the top 4 statistically significant ($P_{adj} < 0.05$) annotations plotted by each gene class (X-axis) as derived from our linear model.

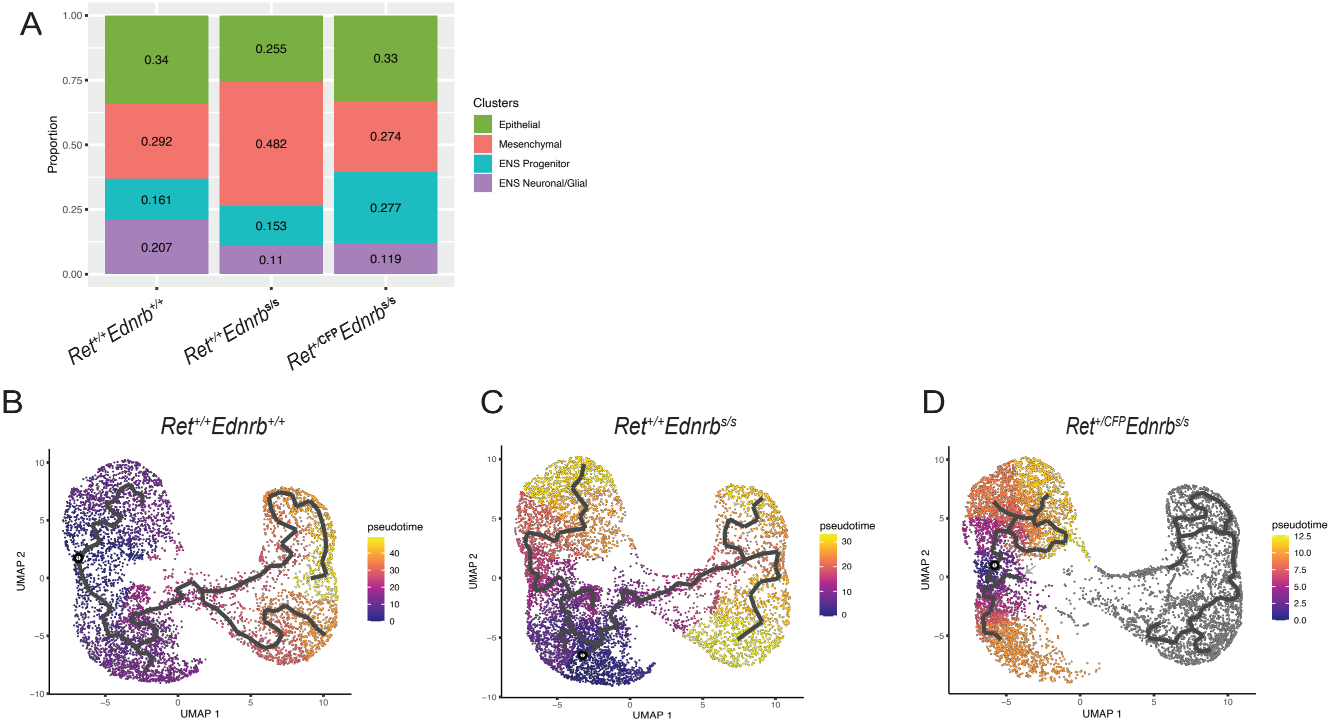


**Figure S6. Epithelial to mesenchymal neuronal trajectory shift in the ENS in *Ret-Ednrb* mutant genotypes (A)** Stacked boxplot of cell type proportions across genotype. **(B)** Pseudotime analysis of WT gut cells reveals a developmental trajectory from epithelial to neuronal cells. **(C)** Analyses as in **(B)** from $Ret^{+/+}$ $Ednrb^{s/s}$ gut cells that shows shift of epithelial cells towards a mesenchymal state (smaller pseudotime distance) between WT and homozygous piebald mice. **(D)** Analyses as in **(B)** from $Ret^{+/CFP}$ $Ednrb^{s/s}$ gut cells reveal a loss of intermediate mesenchymal-like cells and no clear pseudotime path from epithelial to neuronal cell fates (gray arrow).
